# *Escherichia coli* B-strains are intrinsically resistant to colistin and not suitable for characterization and identification of *mcr* genes

**DOI:** 10.1101/2023.02.24.529993

**Authors:** Anna Schumann, Alexa R. Cohn, Ahmed Gaballa, Martin Wiedmann

## Abstract

Antimicrobial resistance is an increasing threat to human and animal health. Due to the rise of multi- and extreme drug resistance, last resort antibiotics, such as colistin, are extremely important in human medicine. While the distribution of colistin resistance genes can be tracked through sequencing methods, phenotypic characterization of putative antimicrobial resistance (AMR) genes is still important to confirm the phenotype conferred by different genes. While heterologous expression of AMR genes (e.g., in *E. coli*) is a common approach, so far, no standard method for heterologous expression and characterization of *mcr* genes exists. *E. coli* B-strains, designed for optimum protein expression, are frequently utilized. Here we report that four *E. coli* B-strains are intrinsically resistant to colistin (MIC 8 – 16 μg/mL). Additionally, the three tested B-strains that encode T7 RNA show growth defects when transformed with empty or gene-expressing pET17b plasmids and grown in the presence of IPTG; K-12 or B-strains without T7 RNA polymerase do not show these growth defects. The *E. coli* Shuffle T7 express strain carrying empty pET17b also showed a heteroresistance phenotype when exposed to colistin in the presence of IPTG. These phenotypes could explain why B-strains were erroneously reported as colistin susceptible. Analysis of existing genome data identified one nonsynonymous change in each *pmrA* and *pmrB* in all four *E. coli* B-strains; the E121K change in PmrB has previously been linked to intrinsic colistin resistance. We conclude that *E. coli* B-strains are not appropriate heterologous expression hosts for identification and characterization of *mcr* genes.

## INTRODUCTION

Antimicrobial resistance (AMR) is an increasing problem and has been declared one of the top 10 global public health threats to humanity by the World Health Organization (WHO) (https://www.who.int/news-room/fact-sheets/detail/antimicrobial-resistance). The Centers for Disease Control and Prevention (CDC) reported that previous gains mitigating the spread of AMR have been reversed due to the COVID-19 pandemic and associated challenges in health care, including a monolithic focus of public health efforts on COVID-19 (1). As AMR spreads and more bacteria become multi- or extremely-drug resistant, our current antibiotics become ineffective and new drugs are needed to combat infections (2)(https://www.who.int/news-room/fact-sheets/detail/antimicrobial-resistance). Unfortunately, new drugs are lacking, so healthcare providers increasingly rely on last-resort antibiotics like colistin (3–6). Colistin, also called polymyxin E, is a cationic antimicrobial peptide discovered in 1947 (6, 7). Even though colistin is nephro- and neurotoxic, colistin use in human medicine has increased in recent years, such that colistin has been defined as a “critically important antimicrobial for human medicine” by WHO (4, 5, 8, 9).

Colistin’s activity is restricted to a few Gram-negative pathogens, including but not limited to members of the order *Enterobacterales* and the species *Pseudomonas aeruginosa*, due to its mechanism of action (5, 6). Colistin has a positively charged head-group, which is electrostatically attracted to the negatively charged phosphate groups of lipopolysaccharide (LPS) molecules within the outer membrane (OM) of Gram-negative bacteria (4, 5). Colistin competitively displaces the Mg^2+^ and Ca^2+^ ions that normally surround the LPS molecules to stabilize the OM (4, 5). The displacement disrupts the OM and causes the leakage of cellular material, lysis, and cell death, although the exact mechanism of cell death is unknown (4, 6, 10). It has been hypothesized that damage to the cytoplasmic membrane may be a key mechanism that contributes to cell death (10).

As with all antibiotics, soon after colistin was used as an antibiotic, resistance was discovered. Originally it was thought that resistance was only conferred by chromosomal mutations in genes regulating expression of LPS modifying enzymes (7, 11). More specifically, modifications made by enzymes EptA and ArnT, which transfer phosphoethanolamine or 4-amino-4-deoxy-L-arabinose, respectively, to the lipid A portion of LPS, can “hide” the negative charge of lipid A, thus blocking colistin’s electrostatic interaction with LPS (7, 11–13). The production of EptA and ArnT is regulated by two separate two-component systems PhoPQ and PmrAB; missense mutations in *phoPQ* and *pmrAB* can cause constitutive activation of lipid-modifying enzymes that result in lipid A modification and thus colistin resistance (7, 12, 14–17). However, in 2015 the first plasmid-encoded gene providing colistin resistance, named mobile colistin resistance (*mcr*) gene, was discovered (18); this gene encodes a protein with a structure similar to EptA. Since then nine more variants (i.e. *mcr-2* – *mcr-10*) and many more sub-variants (e.g. *mcr-2.2*) have been discovered (19–27).

To determine whether a given gene confers colistin resistance and to assess the level of colistin resistance conferred by a given variant or sub-variant, authors oftentimes express *mcr* genes in heterologous laboratory *E. coli* strains using either the native plasmid (18, 28, 29) or cloning the gene into a plasmid downstream of an inducible promoter like the T7 RNA polymerase promoter with the *lac* operator (e.g., pET plasmids) (30, 31). This method is necessary when discovering new variants from whole genome sequence (WGS) data or when the isolate is not available or culturable. However, many different strain and plasmid combinations have been used in the past, making it is difficult to compare resistance levels obtained from different heterologous expression systems (7, 28). The expression host *E. coli* B-strains and their derivatives are frequently used to express recombinant proteins because they were engineered to optimize protein production through their lack of flagella, encoding few proteases, and having increased amino acid biosynthesis (33, 34).

As part of efforts to create a standardized and universal expression system for identification and characterization of *mcr* genes, we noticed that several *E. coli* B-strains appeared to be intrinsically resistant to colistin, despite the fact that *E. coli* B-strains (e.g., BL21(DE3)) have frequently been used to characterize *mcr* genes (30, 31). We thus performed a comprehensive analysis of multiple parent *E. coli* B-strains as well as corresponding strains with the pET17b expression plasmid with and without a cloned *mcr-3*-FLAG. Our data support that *E. coli* B-strains are intrinsically resistant to colistin (with colistin minimum inhibitory concentration [MIC] levels between 8 and 16 μg/mL) with mutations in *pmrAB* representing possible mechanism responsible for the observed resistance phenotype. While overexpression of *mcr-3* using pET17b in B-strains caused a small increase in MIC relative to control strains with an empty plasmid, we also identified substantial toxicity to the *E. coli* host strains that contain pET plasmid in the presence of IPTG. In addition, a specific B strain (i.e., SHuffle T7 express) showed heteroresistance when expressing the empty plasmid. Combined, our data strongly suggest that *E. coli* B-strains, particularly when used with pET plasmids, may not be appropriate to identify and characterize *mcr* genes.

## RESULTS

### Four *E. coli* B-strains show intrinsic resistance to colistin, while two K-12 strains show sensitivity to colistin

During preliminary experiments assessing different host-strain-plasmid combinations for their suitability for heterologous expression of *mcr* genes, the tested B-strains (i.e., BL21(DE3), T7 express lysY/l^q^) showed high levels of resistance to colistin (8-16 μg/mL). For reference, the Clinical and Laboratory Standards Institute (CLSI) and the European Committee on Antimicrobial Susceptibility Testing (EUCAST) define colistin resistance of Enterobacterales as an MIC of or greater than 4 μg/mL (35) or MIC of greater than 2 μg/mL, respectively (36). Based on these findings as well as some previous reports (37, 38), we hypothesized that B-strains are intrinsically resistant to colistin, while K-12 strains are not. To test this hypothesis, we performed MIC assays on four *E. coli* B-strains, including (i) BL21, (ii) BL21(DE3) (abbreviated as “BL21D”), (iii) T7 express lysY/l^q^ (abbreviated as “T7”), and (iv) SHuffle T7 express (abbreviated as “Shuffle”) as well as two *E. coli* K-12 strains, including (i) NEB5α and (ii) Top10; all data were collected in parent strains without expression plasmid grown in media without IPTG. While the four B-strains showed MICs of 8 μg/mL (BL21 and BL21D) and 16 μg/mL (T7 and Shuffle), both K-12 strains showed MICs of <0.5 μg/mL (Table 1).

**Table 1.** MICs of B- and K-12 strains

| Strain | lineage | MIC (μg/mL)<sup>a</sup> |
| --- | --- | --- |
| *E. coli* BL21 | B | 8 |
| *E. coli* BL21(DE3) | B | 8 |
| *E. coli* T7 express lysY/l<sup>q</sup> | B | 16 |
| *E. coli* SHuffle T7 express | B | 16 |
| *E. coli* NEB5α | K-12 | < 0.5 |
| *E. coli* Top10 | K-12 | < 0.5 |

<sup>a</sup>MIC values are based on three biological replicates (except for SHuffle where n=2);
in all cases all replicates yield the same MIC value

### Relative to *E. coli* K-12, *E. coli* B-strains shows nonsynonymous substitutions in *pmrA* and *pmrB*, two genes known to contribute to colistin resistance

Previous papers have reported mutation hotspots in *pmrA* and *pmrB* that have been linked to colistin resistance (16, 37). We thus hypothesized that the increased colistin MIC values observed here in the four *E. coli* B-strains, as compared to the two K-12 strains, could be due to mutations in *pmrA* and/or *pmrB*. To test this hypothesis, we aligned the *pmrA* and *pmrB* genes of the four B-strains and K-12 strain NEB5α to the *pmrA* and *pmrB* genes of *E. coli* K-12 MG1655 (as a reference strain); *E. coli* Top10 *pmrA* and *pmrB* are not included in this alignment as we were unable to obtain a Top10 genome sequence. The *pmrA* alignment identified a total of 7 synonymous and one nonsynonymous site over the 669 nucleotides (nt) length of the coding sequence (Fig. S1). For all 8 sites, all four *E. coli* B-strains showed the same sequences and both NEB5α and MG1655 showed the same sequences, consistent with their classification into different *E. coli* lineages (i.e., B and K-12). For the nonsynonymous substitution at amino acid site 29, the four B-strains encoded a glycine, while the two K-12 strains encoded a serine (Fig. 1). The *pmrB* alignment identified one synonymous and one nonsynonymous site over the 1,092 nt length of the coding sequence (Fig. 1). Again, for the nonsynonymous site, all four *E. coli* B-strains showed the same sequence and both K-12 strains showed the same sequences, while for the synonymous substitution NEB5α was the only strain that deviated from the consensus DNA sequence (Fig. 1). For the nonsynonymous substitution at amino acid site 121, the four B-strains encode a lysine, while the two K-12 strains encode a glutamate (Fig. 1).

**Figure 1.**
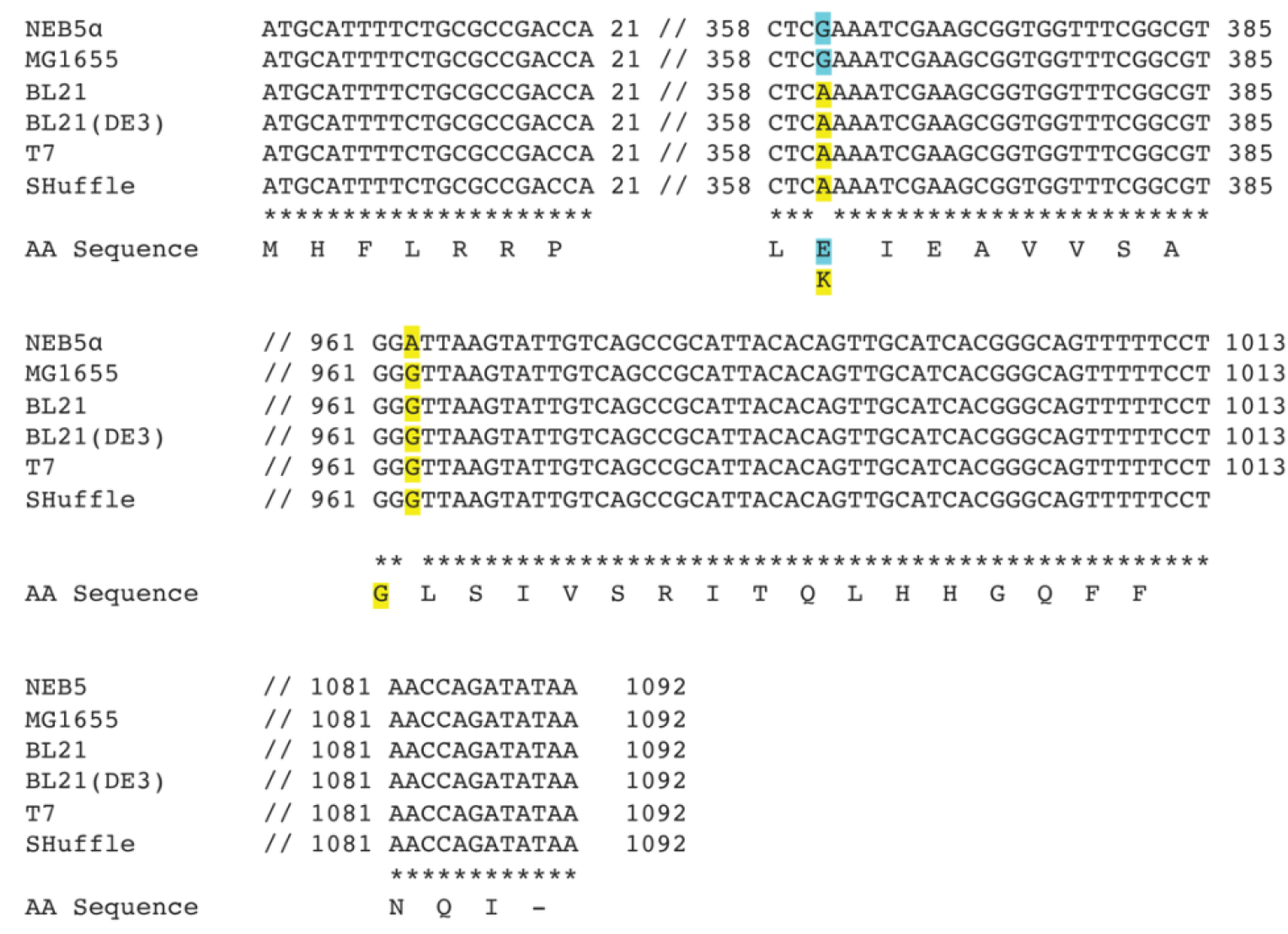
Alignment of *pmrB* DNA sequences of *E. coli* B- and K-12 strains used in this study. Alignment of genes and matching amino acid sequences was performed by Clustal Omega (48). Sequences from nucleotides 22 – 357, 386 – 960, and 1014 – 1080 were omitted because the strains shared 100% sequence identity for each of these regions. Base changes are highlighted; coloring indicates which base change correlates with which amino acid. Refer to materials & methods section for details on how the sequences were acquired. [T7 = T7 express lysY/l^q^, SHuffle = SHuffle T7 express]

### IPTG-induced *E. coli* B-strains carrying pET17b-mcr-3-FLAG plasmid show 2-fold higher colistin MIC values as compared to empty plasmid controls without pET17b

As B-strains have been used in a few studies to characterize *mcr* genes, despite evidence for intrinsic colistin resistance, we generated three different lineage B-strains (i.e., BL21D, T7, Shuffle) that each either carry a pET17b plasmid expressing *mcr-3* (pET17b-mcr-3-FLAG) or an empty pET17b plasmid. We chose to use *mcr-3* because the colistin MIC values of heterologous strains expressing *mcr-3* are well characterized and should be 2-4 μg/mL, depending on the heterologous expression system used and *mcr-3* subvariant (25, 39, 40). Colistin MIC determinations of these strains under uninduced conditions (0 mM IPTG) as well as under induction (0.4 and 1 mM IPTG) showed 2-fold higher MIC values for induced strains expressing *mcr* as compared to the corresponding strains carrying an empty plasmid (Table 2). For example, BL21D expressing *mcr-3* showed MIC values of 8 μg/mL (for both 0.4 and 1 mM IPTG), while the same strain with an empty plasmid showed an MIC of 4 μg/mL at both IPTG concentrations. By comparison, when strains were not induced with IPTG, the *E. coli* strains expressing *mcr* showed the same MIC values as the strains carrying an empty plasmid; for example, BL21D strains with the pET17b-*mcr-3* and with the empty plasmid both showed MIC values of 4 μg/mL (Table 2). We confirmed MCR-3-FLAG protein expression by western blot analysis after the induction of expression in B-strains BL21D, T7, and Shuffle with 0.4 mM IPTG and dH2O as a negative control (Fig. S2). As expected, we did not observe expression of MCR-3-FLAG in the strains carrying pET17b but observed it in all strains carrying pET17b-*mcr-3* (Fig. S2).

**Table 2.** MICs of *E. coli* B-strains carrying pET17b (“EV”) and pET17b_*mcr-3*-FLAG (“mcr-3”) after induction with different IPTG concentrations

| strain | plasmid | MIC (µg/mL) obtained with<sup>a</sup> |  |  |
| --- | --- | --- | --- | --- |
|  |  | 0 mM IPTG | 0.4 mM IPTG | 1 mM IPTG |
| BL21 (DE3) | No plasmid | 8 | - | - |
|  | EV | 4 | 4 | 4 |
|  | *mcr*-3 | 4 | 8 | 8 |
| T7 express<br>lysY/l<sup>q</sup> | No plasmid | 16 | - | - |
|  | EV | 16 | 8 | 8 |
|  | *mcr*-3 | 16 | 16 | 16 |
| SHuffle T7<br>express | No plasmid | 16 | - | - |
|  | EV | 16 | 0.5/8<sup>b</sup> | 0.5/16<sup>c</sup> |
|  | *mcr*-3 | 16 | 16 | 16 |

<sup>a</sup>MIC values represent the majority of three biological replicates; - indicates samples for which no MIC data were collected;
<sup>b</sup>One replicate each skipped one (0.5 µg/mL) or two wells (0.5 and 1 µg/mL);
<sup>c</sup>All three replicates skipped two wells (0.5 and 1 µg/mL)

In addition to the observations detailed above, we also found that for two of the three B-strains (i.e., T7 and Shuffle), the MIC for colistin of the strains carrying pET17b-*mcr-3* was the same regardless of IPTG induction (Table 2). For example, T7 pET17b-*mcr-3* showed MIC values of 16 μg/mL when the strain was uninduced as well as when *mcr* expression was induced with 0.4 and 1 mM IPTG (Table 2). When examining the western blot analysis mentioned above, our data also showed that *mcr-3* was expressed even in the absence of IPTG; semi-quantitative analysis of *mcr-3* levels shows that relative *mcr-3* levels in strains induced with IPTG only showed a small increase in *mcr-3* levels as compared to strains grown without IPTG (Fig. S2). These data suggest that leaky expression of *mcr-3*-FLAG occurs and is likely responsible for our findings that colistin MIC values tend to not differ between B-strains that carry pET17b-*mcr-3* regardless of whether expression is induced by IPTG.

### *E. coli* Shuffle carrying pET17b empty plasmid shows heteroresistance

As part of the colistin MIC determination experiments detailed in the preceding section, we also found evidence that at least one B-strain (i.e., Shuffle) can show a heteroresistance phenotype, which is defined as different subpopulations of a strain having different resistance levels to an antibiotic (41). This phenomenon can be observed during standard broth microdilution MIC assays, when strains “skip wells”, e.g., do not show growth at one concentration (e.g., 0.5 μg/mL), but then show growth again at a higher concentration (e.g., 1, 2 and 4 μg/mL) (41). More specifically, we observed a heteroresistance phenotype in broth microdilution MIC assays with the Shuffle strain containing the empty vector (EV) induced with either 0.4 or 1 mM of IPTG. The heteroresistance phenotype, was more prominent with 1 mM IPTG, as all three biological replicates skipped two wells (i.e., 0.5 and 1 μg/mL) but continued to grow at 2, 4, and 8 μg/mL (Table 2), while for 0.4 mM, our replicates skipped one (i.e., 0.5 μg/mL) or two wells (i.e., 0.5 and 1 μg/mL), but all grew at 2 and 4 μg/mL. This observation is important as misinterpretation of heteroresistance (and “skipped wells”) may provide inaccurate MIC values that could ultimately be mis-interpreted as colistin resistance.

### *E. coli* B-strains encoding T7 RNA polymerase and transformed with pET17b show reduced cell viability in the presence of IPTG

In addition to heteroresistance, cytotoxicity of either plasmid constructs and/or specific cloned genes (e.g., *mcr*) can also yield misleading data that can provide inaccurate MIC values and hence lead to incorrect conclusions on resistance phenotypes conferred by a given overexpressed gene (42, 43). As previous reports indicated that over expression of *mcr* genes may be toxic to the *E. coli* host cells (42, 43), we performed experiments to determine whether T7 RNA polymerase encoding *E. coli* B-strains transformed with pET17b-*mcr-3* show reduced cell viability. For these experiments, B-strains without plasmid (“no plasmid”), with pET17b, and with pET17b-mcr-3 were plated on LB agar supplemented with 0, 0.4 and 1 mM of IPTG (Fig. 2). On LB that was not supplemented with IPTG, growth patterns were indistinguishable for each given B-strain, no matter if they carried no plasmid, pET17b, or pET17b-*mcr-3* (Fig. 2). On the other hand, on LB supplemented with 0.4 or 1 mM of IPTG, both BL21D and T7 carrying pET17b and BL21D and T7 carrying pET17b-*mcr-3* showed decreased cell numbers, as compared to the control without IPTG (Fig. 2). Interestingly, for *E. coli* Shuffle, the strain carrying pET17b also showed reduced growth, as compared to the control without IPTG, while the strain carrying pET17b-*mcr-3* did not show reduced cell viability relative to the control without IPTG. In addition, T7 and Shuffle strains carrying pET17b show different colony morphology on LB with 0.4 or 1 mM IPTG, as compared to the LB control without IPTG; in the presence of IPTG these strains predominantly show tiny colonies (as compared to colony sizes observed in the absence of IPTG) with only a few colonies showing a size similar to the colony sizes observed when these strains were grown on LB without IPTG (Fig. 2). Colony size is inversely proportional to toxicity, making it an additional indicator of toxicity (44). As reduced cell viability is observed in strains that are exposed to IPTG and carry an empty plasmid, this phenotype clearly does not solely reflect toxicity associated with *mcr* over-expression.

**Figure 2.**
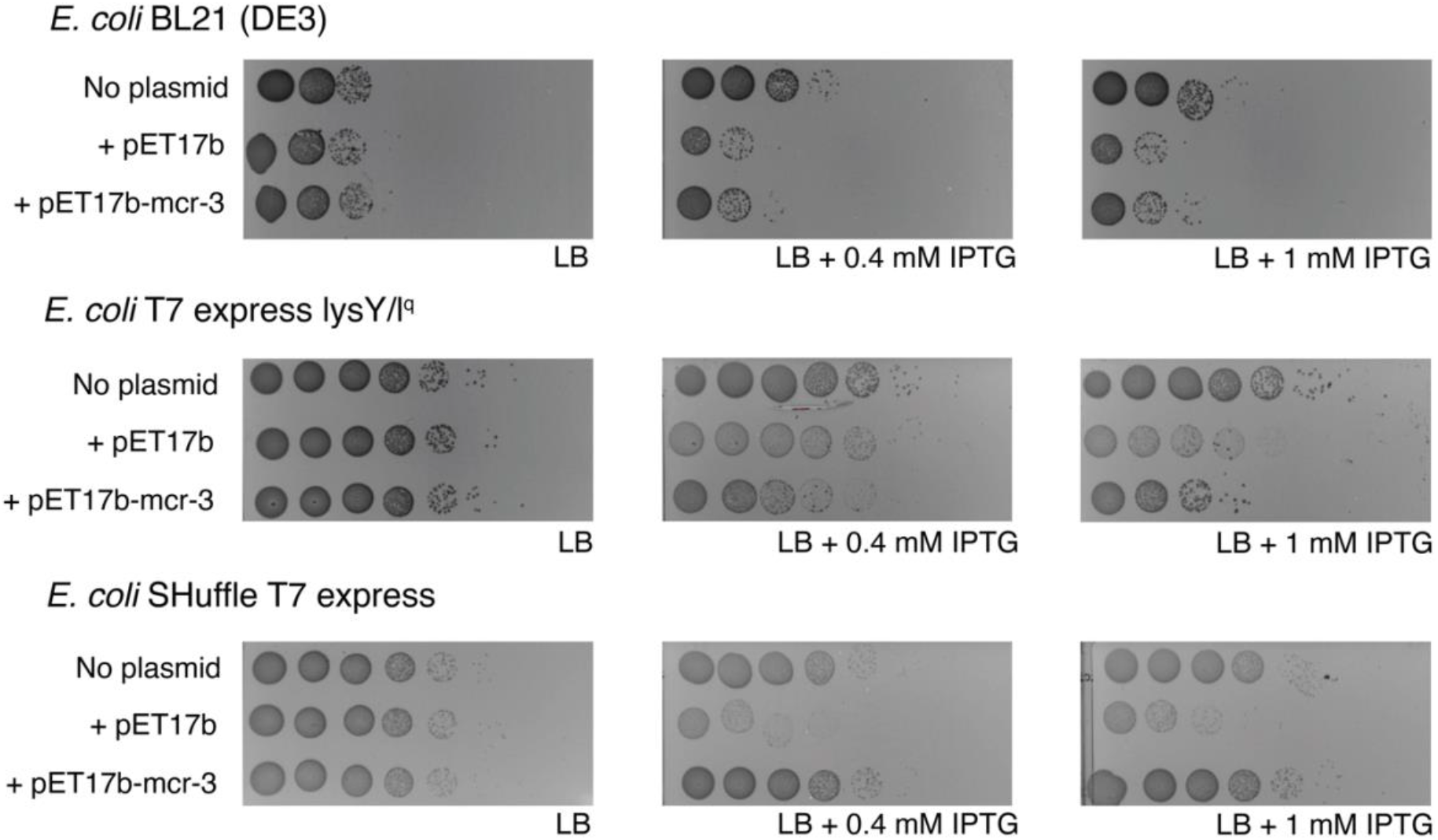
Cell viability of *E. coli* B-strains encoding T7 RNA polymerase on LB agar containing IPTG. Plating assay of *E. coli* B-strains without plasmid (“no plasmid”), with empty pET17b (“+ pET17b”), and with pET17b-*mcr3* (“+ pETI7b-mcr-3”) on 0, 0.4, or 1 mM of IPTG (0mM IPTG plates were prepared by adding distilled water instead of IPTG). Exponentially grown strains were 10-fold serial diluted in PBS. Undiluted to 10^-7^ dilutions were plated in 10μL volumes. Results were consistent across three replicates and one replicate is shown here; images were acquired with the Bio-Rad ChemiDoc MP Imaging system.

To determine whether the growth defect of B-strains that carry pET17b – either empty or expressing *mcr-3* – supplemented with IPTG is due to plasmid toxicity or due to combined effect of the plasmid and T7 RNA polymerase expression, we transformed pET17b and pET17b-*mcr-3* into BL21 and NEB5α, a B and K-12 strain, respectively. These two strains do not carry a copy of T7 RNA polymerase and hence are not capable of using the T7 promoter that controls expression of recombinant genes in pET17b. When we plated BL21 and NEB5α (each without plasmid, with pET17b and pET17b-*mcr-3*) on LB agar with 0, 0.4, and 1 mM of IPTG, bacterial numbers (as well as colony morphology) were indistinguishable in the presence and absence of IPTG regardless of whether a given strain carried no plasmid, pET17b, or pET17b-*mcr-3* (Fig. 3).

**Figure 3.**
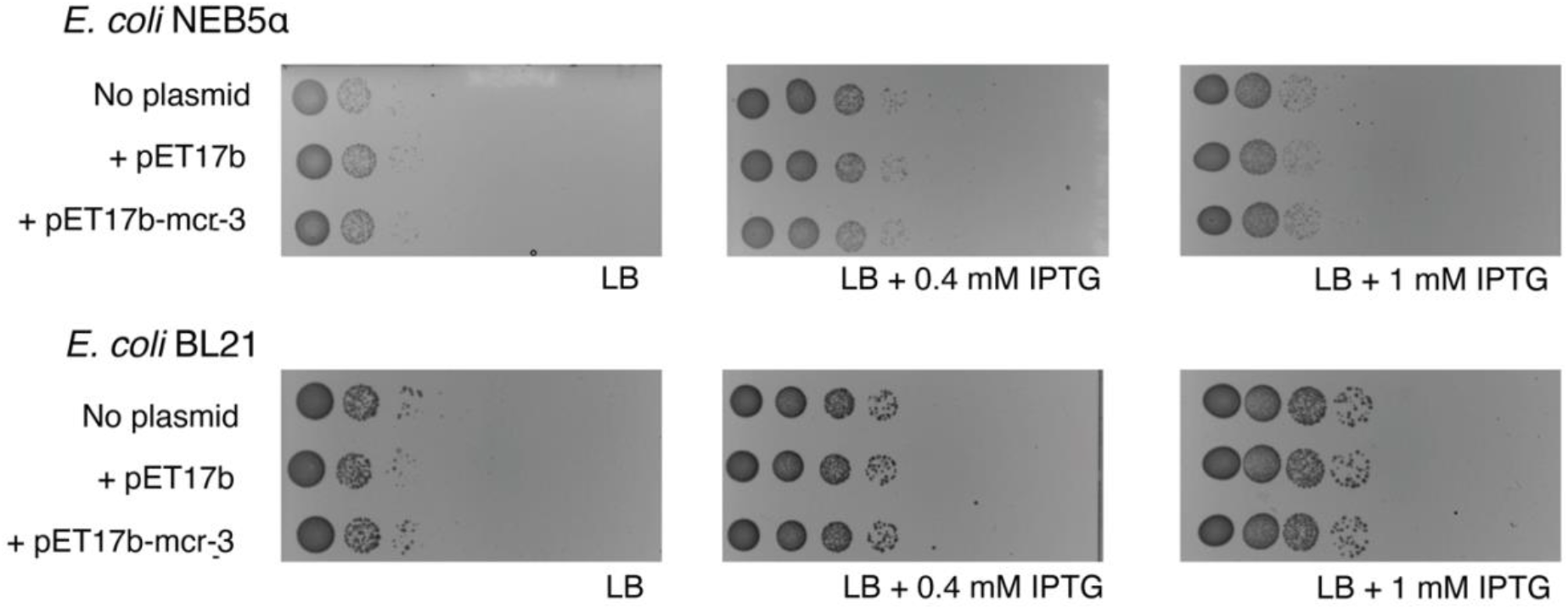
Cell viability of *E. coli* strains without T7 RNA polymerase on LB agar containing IPTG. Plating assay of *E. coli* strains without T7 RNA polymerase without plasmid (“no plasmid”), with empty pET17b (“+ pET17b”), and with pET17b-*mcr-3* (“+ pET17b-mcr-3”) on 0, 0.4, or 1 mM of IPTG (0mM IPTG plates were prepared by adding distilled water instead of IPTG). Exponentially grown strains were 10-fold serial diluted in PBS. Undiluted to 10^-7^ dilutions were plated in 10μL volumes. Results were consistent across three replicates and one replicate is shown here; images were acquired with the Bio-Rad ChemiDoc MP Imaging system.

As the toxicity seems limited to strains carrying T7 RNA polymerase, we hypothesized that T7 RNA polymerase expression may cause metabolic burden or interfere with specific cellular processes or cellular homeostasis. It has been previously shown that overexpression of T7 RNA polymerase may reduce magnesium availability, as it uses magnesium as a cofactor (45). We thus used LB agar supplemented with 1 mM of MgCl_2_ to repeat cell viability experiments with two B-strains encoding T7 RNA polymerase (i.e., BL21D; T7) with either no plasmid, pET17b, or pET17b-*mcr-3* in the presence of 0, 0.4, and 1 mM of IPTG. Our results show that MgCl_2_ does not appear to alleviate the cell viability effects seen when adding IPTG to pET17b carrying B-strains (Fig. 4), as we see reduced bacterial numbers for both BL21D and T7 (carrying pET17b or pET17b-mcr-3), similar to the patterns observed in LB not supplemented with MgCl_2_ (Fig. 4). Similarly, T7 carrying pET17b also showed the same atypical colony morphology (i.e., predominantly tiny colonies) on LB supplemented with MgCl_2_ as was observed on LB not supplemented with MgCl_2_. Our results do not support our hypothesis, meaning that the decrease in cell numbers is not associated with reduced availability of cellular magnesium, but rather suggest that cytotoxicity may be associated with the metabolic burden caused by the overexpression of T7 RNA polymerase.

**Figure 4.**
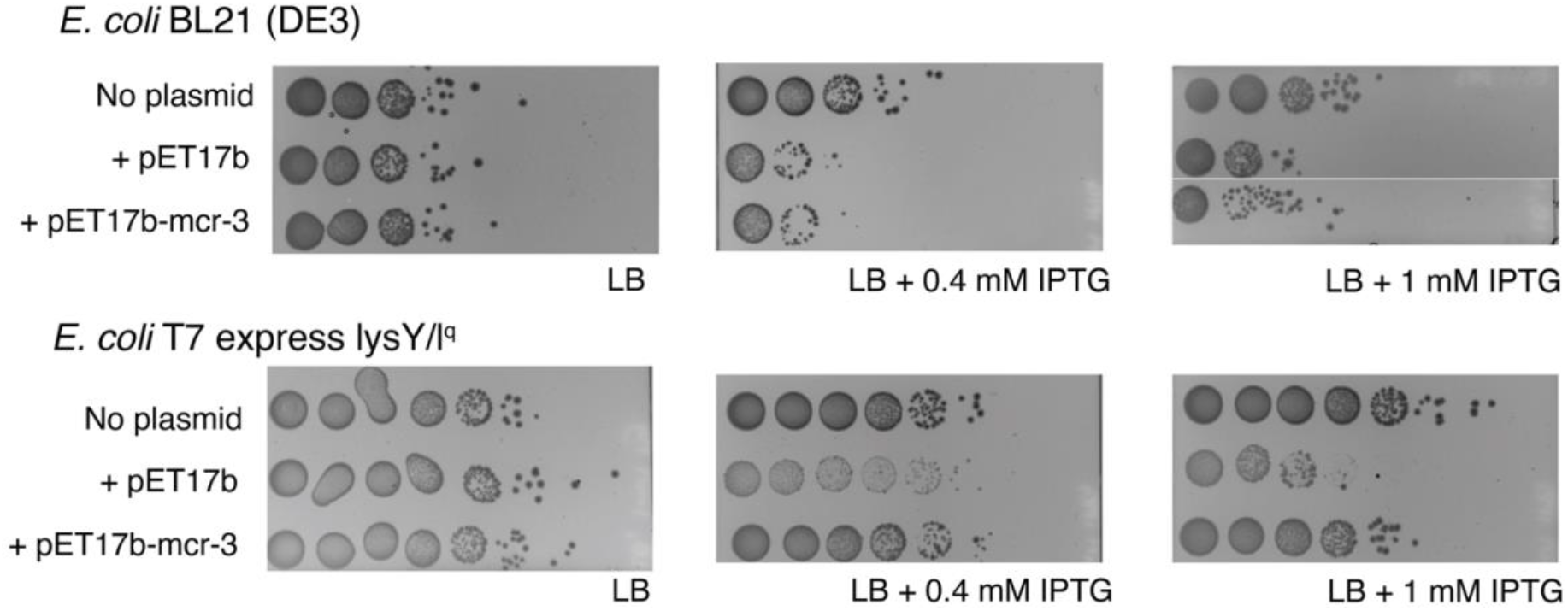
Cell viability of *E. coli* BL21D and T7 on LB agar containing 1 mM MgCl_2_ and IPTG. Plating assay of exponentially growing *E. coli* B strains BL21D and T7 without plasmid (“no plasmid”), with empty pET17b (“+ pET17b”), and with pET17b - mcr-3 (“+ pET17b-mc-r3”) on LB containing 1 mM MgCl_2_ and 0, 0.4, or 1 mM of IPTG (0mM IPTG plates were prepared by adding distilled water instead of IPTG). Exponentially grown strains were 10-fold serial diluted in PBS. Undiluted to 10^-7^ dilutions were plated in 10μL volumes. Results were consistent across three replicates and one replicate is shown here; images were acquired with the Bio-Rad ChemiDoc MP Imaging system.

## DISCUSSION

While antibiotic resistance determination (e.g., MIC determination) on clinical and other wildtype strains remains a cornerstone of antimicrobial resistance surveillance, heterologous expression of AMR genes represents another important approach for AMR resistance characterization. Advantages and applications of heterologous expression of AMR genes include (i) ability to characterize resistance conferred independent of genetic background (which may affect resistance gene expression, e.g., if AMR genes expression is tightly regulated and does not occur in wildtype strains grown under lab testing conditions), and (ii) ability to characterize resistance phenotypes if wildtype strains are not available (e.g., through using synthetic resistance genes). For example, in some cases genome sequences may indicate presence of putative novel AMR genes, but the actual strains may not be available to researchers. With widespread global use of Culture-independent diagnostic techniques (CIDT) and WGS technologies (and a corresponding decrease of routine wildtype strain MIC determinations), use of novel approaches to identify and characterize emerging AMR genes (e.g., synthesis of genes found in genome databases, followed by characterization in heterologous expression hosts) is becoming more important. Importantly, similar approaches are already being used, including for the characterization of *mcr* genes (23, 25, 29, 42, 46). A lack of standardization of these methods, including the use of different heterologous expression hosts (which often were optimized for maximum protein expression) however represents a major challenge. Here we show that (i) commonly used heterologous expression hosts (i.e., *E. coli* B-strains) show intrinsic resistance to colistin, and (ii) commonly used overexpression systems (i.e., IPTG induction of pET17b plasmids in B-strains encoding T7 RNA polymerase) can induce cytotoxicity and other aberrant phenotypes (i.e., heteroresistance), which is expected to lead to erroneous MIC results. While our findings indicate that *E. coli* B-strains should not be used as hosts for *mcr* variant MIC determinations, we also showed that *E. coli* K-12 strains appear to be appropriate hosts for characterization and identification of *mcr* genes. More broadly, our findings suggest that improved evaluation and standardization of heterologous expression system-based AMR determinations is urgently needed, particularly as use of CIDT based diagnostics increases.

### Commonly used heterologous expression hosts (i.e., *E. coli* B-strains) show intrinsic resistance to colistin, likely aided by mutations in *pmrAB*

We found that four *E.coli* B-strains, BL21, BL21D, T7, and Shuffle are intrinsically resistant to colistin with MICs of 8-16 μg/mL. This observation was surprising, as previous studies have used B-strains like BL21D and Shuffle to report MIC values for *mcr* variants and indicated that the strains were susceptible to colistin (26, 30, 31). However, our findings are consistent with studies by Trent et al. (38) and Xu et al. (37), who reported that B-strains BLR(DE3) (obtained from Novagen) and BL21(DE3) (obtained from both Novagen and Stratagene), used in their respective studies, were resistant to colistin. Trent et al. (38) attributed the colistin resistance of BLR(DE3) to phosphoethanolamine and L-arabinose modifications of the lipid A portion of LPS as supported by the observation, based on analyzing ^32^P-labeled lipid A species, that colistin resistant *Salmonella* Typhimurium *pmrA* mutant and *E. coli* BLR(DE3) showed similar lipid A modifications, which were different from colistin susceptible *E. coli* K-12 strain NovaBlue(DE3) (38). Overall, our data combined with these previous findings suggest that B-strains are not suited as heterologous expression hosts for characterization and identification of *mcr* genes. While prior evidence of colistin resistance in *E. coli* B-strains has been reported by Trent et al. (38) in 2001, this previous paper had focused on characterization of aminoarabinose transferase (ArnT) and thus appears to have been largely ignored as a substantial number of subsequent studies (26, 30, 31) still used *E. coli* B-strains for heterologous expression of *mcr* in order to identify and characterize *mcr* genes.

Comparative genomic analysis of existing genomes of the four *E. coli* B-strains used here, as well as two K-12 strains, NEB5α and MG1655, identified one nonsynonymous change in each *pmrA* and *pmrB*. Specifically, B-strains encode a glycine at amino acid site 29 in PmrA and a lysine at amino acid site 121 in PmrB, while K-12 strains encode a serine and a glutamate, respectively, at these locations. Mutations in the two-component system PmrAB represent a plausible mechanism for colistin resistance, as PmrAB regulates the expression of lipid A modification system, and many studies have described mutations in *pmrAB* that impact colistin resistance (8, 13–16, 37, 47). The serine to glycine change in PmrA has also previously been identified in two colistin resistant *E. coli* isolates from chicken feces (14). These isolates also had one nonsynonymous mutation in *pmrB* and the authors did not experimentally confirm whether the S29G change in PmrA conferred resistance on its own (14). Because serine and glycine have weakly similar properties according to Clustal Omega (48), it is unclear whether this change actually affects the expression of the lipid A modification system and, thus, colistin resistance. On the other hand, the glutamate to lysine change in PmrB has been identified in multiple studies with different conclusions as to whether this change confers colistin resistance or not (16, 37, 47). While one study (16) identified the E121K change in more than ten isolates, they did not experimentally test whether this PmrB mutation conferred resistance. Another study (47) classified this mutation as deleterious, using the PROVEAN tool, which predicts the effect of amino acid substitutions on protein function (49). Xu et al. identified the same change in PmrB in laboratory *E. coli* BL21(DE3) strains acquired from Stratagene and Novagene (37) and showed that colistin resistance increased when the *pmrB* with at least 126 bp of the 3’ untranslated region (UTR) was expressed in an *E. coli* Top10 strain. However, cloning of *pmrB* without the 3’ UTR or with a shorter 3’ UTR did not appear to confer colistin resistance, suggesting a possible contributing role of mRNA stability. Expression of *pmrB* with the 126 bp 3’UTR also resulted in phosphoethanolamine and L-arabinose modifications of lipid A (37), which could be linked to the observed colistin resistance. Overall, previous data support that mutations in both PmrA and PmrB can confer colistin resistance. While it thus is plausible that the combination of amino acid changes in PmrA and PmrB is the reason why B-strains are resistant to colistin, additional experimental work would be necessary to prove causality. We however do not see these types of mechanistic studies as necessary for this paper, as our data, along with previous studies, sufficiently demonstrate that B-strains are resistant to colistin and should therefore not be used to identify and characterize *mcr* genes.

### Commonly used overexpression systems can induce cytotoxicity and other aberrant phenotypes, which is expected to lead to erroneous colistin MIC results

Our study found evidence that induction of T7 RNA polymerase expression induces toxicity in strains that carry the expression plasmid pET17b without an insert. This was supported by the finding that B-strains that encode T7 RNA polymerase and carry the empty plasmid showed reduced growth in the presence of IPTG, as compared to growth in media without IPTG, while strains that do not encode T7 RNA polymerase (i.e., BL21, NEB5α) did not show any differences in cell growth between media with and without IPTG. Our findings are consistent with a study (50) that reported a growth defect in *E. coli* BL21 (DE3) carrying an empty plasmid, when induced with IPTG. The authors attributed the growth defect in those strains to the increased metabolic burden on the cells due to expression of selective marker and maintenance of the plasmid (50). Importantly, in BL21D, T7, and Shuffle, expression of T7 RNA polymerase is controlled through the addition of IPTG (https://www.neb.com/products/competent-cells/e-coli-expression-strains/e-coli-expression-strains) and induction of T7 RNA polymerase expression in strains carrying the empty pET17b plasmid has been reported to initiate transcription from the T7 promoter upstream of the multiple cloning site (MCS) even if no gene is cloned into the MCS (https://research.fredhutch.org/content/dam/stripe/hahn/methods/biochem/pet.pdf).

While this transcription initiation would be expected to create a short transcript (and not a translated protein), it has been shown that the native T7 terminator in pET17b is not 100% efficient, which can lead to read-through transcription (51, 52). If read-through transcription occurs, the downstream ampicillin resistance gene, which encodes a β-lactamase, will be transcribed, translated, and secreted (33, 53). Previous studies have shown that the overexpression of secreted and membrane proteins can result in toxicity due to the formation of cytoplasmic aggregates because the pathways for protein production and secretion are saturated (54, 55). The increased burden of gene expression and saturation of the secretion machineries through read-through transcription could thus explain the toxicity we and others have seen in IPTG-induced B-strains encoding T7 RNA polymerase (e.g., BL21(DE3)) carrying the empty pET17b plasmid.

Our data also support that the observed cytotoxicity in T7 RNA polymerase-expressing strains can provide aberrant colistin MIC patterns (even in empty pET17b carrying strains), which could be misinterpreted as the strain being susceptibility to colistin. Specifically, we found that for both BL21D and T7, strains that carry pET17b-*mcr-3* that were induced with IPTG show the same colistin MIC as the parent strain without plasmid, while showing a 2-fold higher MICs as compared to control strains that carried an empty pET17b and were induced with IPTG. Hence, strain expressing *mcr-3* would be considered as showing evidence for “resistance” if compared to an IPTG induced strain with an empty plasmid (possibly due to cytotoxicity displayed by this strain) but would not show evidence for resistance when compared to the parent strain without a plasmid. Importantly, we also found evidence for toxicity in *E. coli* BL21D and T7 that carry pET17b-*mcr-3* when T7 RNA polymerase expression is induced by IPTG. This observation is consistent with previous studies showing that overexpression of heterologous *mcr* genes in lab strains causes cytotoxicity, presumably because MCR and/or MCR induced lipid A modifications disrupt the bacterial cell membranes and impact membrane stability (42, 43). As our western blots showed that MCR-3-FLAG is expressed even in the absence of IPTG induction, most likely due to the leakiness of the *lac* promoter that controls expression of T7 RNA polymerase which in turn expresses *mcr-3* (56, 57), our results suggest that *mcr-3* expression is only toxic in BL21D and T7 when expression occurs at higher levels. Overall, these data show that a number of potentially direct and indirect factors affecting cytotoxicity phenotypes can occur when using T7 RNAP encoding parent strains and pET overexpression plasmids to phenotypically assess different *mcr* genes.

In addition to the cytotoxicity phenotypes detailed above, we also found that Shuffle strains carrying the empty plasmid showed a heteroresistance phenotype, which was not observed with BL21D and T7. Heteroresistance has been reported in 1947 for the first time and can display as “skipping wells” in broth microdilution (BMD) MIC assays (41). This phenomenon of “skipped wells” is known to occur in *Enterobacter* spp. in response to colistin (58, 59) and has also been reported recently by Kananizadeh et al. (60) when testing wildtype isolates carrying *mcr-9* in MHII broth supplemented with peptone, tryptone or casein. To our knowledge we are the first to report the phenomenon when testing *mcr* genes or their empty expression plasmids in heterologous expression strains grown in standard MHII broth. According to CLSI guidelines, MICs of strains that skip two or more wells in BMD assays should not be reported (61), however, it is plausible that skipped wells may be misreported as a lower MIC than the “true MIC”. For example, one may interpret growth at a higher colistin concentration (e.g., 4 μg/mL) as an “outlier” or “contamination event” and discount that data point and hence interpret the skipped wells at 1 and 2 μg/mL as inhibition, thus assigning an MIC of 0.5 μg/mL, when the true MIC value is 4 μg/mL. This is a particular concern and challenge if heteroresistance is only observed with an empty plasmid, as this may lead to an incorrectly assigned low MIC value of a negative control strain. Comparison of this “incorrectly low” MIC value to the MIC value in a strain expressing a resistance gene (e.g., *mcr-3*) could thus misclassify the strain expressing a resistance gene as showing a higher MIC, and hence, resistance. This possibility is illustrated with our data for the Shuffle strain; here, incorrect interpretation of heteroresistance would yield an MIC value of 0.5 μg/mL for the Shuffle strain with the empty plasmid, while the Shuffle strain with pET17b-*mcr-3* showed an MIC value of 16 μg/mL which overall would be mid-interpreted as *mcr-3* providing colistin resistance in this strain.

Based on the cytotoxicity phenotypes observed with both IPTG induction of B-strains encoding T7 RNA polymerase carrying the empty pET17b plasmid and with overexpression of *mcr* as well the heteroresistance (“skipped well”) phenomenon observed for Shuffle strains carrying pET17b, we suggest that the commonly used expression system of B-strains with T7 RNA polymerase and pET plasmids may not be appropriate to phenotypically characterize *mcr* genes (and possibly any antimicrobial resistance genes). Specifically, these aberrant phenotypes could lead to misinterpretation of colistin MIC values, particularly when using EUCAST and CLSI cut-off values to determine resistance (35, 36). However, our experiments cannot explain why other studies have reported that the parental B-strains that carry T7 RNA polymerase but do not carry plasmids are susceptible to colistin (30, 31); for example a study (30) has reported a colistin MIC value of 0.5 μg/mL for *E. coli* SHuffle T7 express, while we found a MIC value of 16 μg/mL. Among other reasons, this could be caused by colistin preparations with reduced efficacy, higher strain concentrations than used here and recommended by CLSI (61), pre-growth of strain under non-standard conditions, and possibly even use of parent strains that did not have the colistin resistance phenotype reported here (e.g., due to mutations, contamination with other *E. coli* strains, or sharing of mislabelled or misassigned strains between labs).

### While *E. coli* B-strains should not be used as hosts for colistin MIC determinations, *E. coli* K-12 strains appear to be appropriate hosts for characterization and identification of *mcr* genes

We found that two *E. coli* K-12 strains, NEB5α and Top10, were susceptible to colistin with MICs of <0.5 μg/mL. Our findings of K-12 strain’s MIC values are consistent with a previous study that reported that K-12 strain NovaBlue(DE3) is susceptible to colistin and has no lipid A modifications (38). Other studies have used *E. coli* DH5α (19, 21, 27, 62) – the NEB5α strain that was used here is a derivative of DH5α according to NEB (https://www.neb.com/products/c2987-neb-5-alpha-competent-e-coli-high-efficiency#Product%20Information) - and *E. coli* Top10 as heterologous expression strains for *mcr* variants and reported the colistin MICs of their heterologous expression strains to be similar to ours (28, 63, 64). The levels of colistin resistance are consistent with our genotypic findings of only one synonymous base change in *pmrB* and no change in *pmrA* of NEB5α in comparison to *E. coli* MG1655. Based on our phenotypic and genotypic observations that are supported by previous studies, *E. coli* K-12 strains appear to be appropriate heterologous expression strains for the characterization and identification of *mcr* variants. However, our findings also suggest that it is essential to perform an initial MIC screen with strains and vectors that are being considered as heterologous expression hosts, regardless of which antibiotic resistance gene is being characterized. Based on previous studies, expression plasmids pCR (28), pBADb (63), pUC19 (19), and the native plasmid (27) are some of the plasmids that have been used in conjunction with *E. coli* K-12 strains, DH5α and Top10 to identify and characterize novel *mcr* genes and variants and could provide alternatives to pET plasmids. As *E. coli* K-12 strains DH5α and Top10 do not encode T7 RNA polymerase these strains cannot be used in combination with pET plasmids. When choosing a combination of expression strain and plasmid, it is also important to confirm that the heterologous expression strain encodes the necessary genes for gene expression from the promoter present in the expression plasmid.

Antimicrobial resistance is major threat to human and animal health. Especially when CIDT-based diagnostics are used, heterologous expression systems are essential for antibiotic resistance determination. We have shown that it is important to thoroughly examine whether a heterologous expression strain is compatible with the AMR gene that is being examined, as *E. coli* B-strains that have frequently been used to identify and characterize *mcr* variants are intrinsically resistant to colistin. Additionally, expression systems can result in erroneous determination of MIC values through cytotoxicity or heteroresistance phenotypes as seen here, suggesting that the combinations of expression strains and plasmids should be examined for unexpected phenotypes. Finally, our findings suggest that improved evaluation and standardization of heterologous expression system-based AMR determinations are urgently needed.

## MATERIALS AND METHODS

### Strains, plasmids, and growth conditions

The strains and plasmids used in this study are shown in Table 3. All *S. enterica* strains and *E. coli* strains were grown at 37°C with shaking at 200 rpm, in Difco LB Lennox Broth (LB; Becton, Dickinson and Company [BD]; Franklin Lakes, NJ; cat. #240230). For *E. coli* strains carrying the pET17b plasmids, 100 μg/mL of ampicillin (AMP) was added to maintain the plasmid. All plasmids were transformed into competent *E. coli* strains by heat shock, according to the manufacturer’s recommendations (New England Biolabs [NEB]; Ipswich, MA).

**Table 3.**
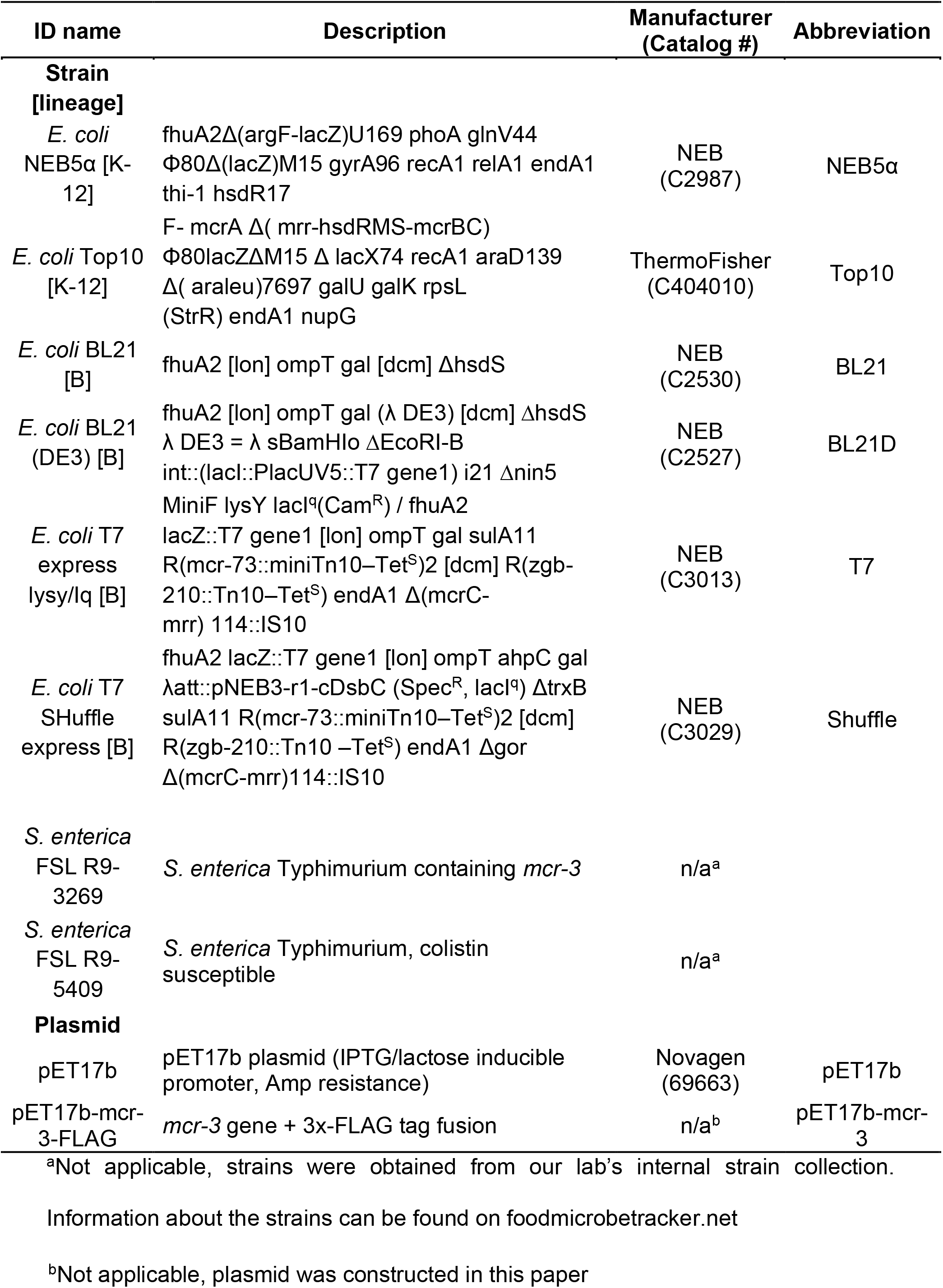
Strains and Plasmids used in the study

### Cloning of *mcr-3*

mcr-3 was amplified via PCR with the primers pET17b_mcr3_F and pET17b_mcr3_R (Table 4) from the genomic DNA of isolate *S. enterica* FSL R9-3269. The PCR amplicon was purified and used as a template to add the 3x FLAG tag through primer walking using the primers pET17b_mcr3_F and pET17b_3x_FLAG_R2 (Table 4). All PCR reactions were performed using Q5 high-fidelity DNA polymerase (NEB, cat. #M0491S) according to the manufacturer’s recommendations. PCR products and the pET17b plasmid (Novagen (EMD Millipore), cat. #69663) were digested with restriction enzymes *NdeI* (NEB, cat. #R0111S) and *Bam*HI (NEB, cat. #R0136S), purified, and ligated with T4 DNA ligase (NEB, cat. #M0202L) according to the manufacturer’s recommendations. Ligation products were transformed via heat shock into *E. coli* Lemo21(DE3) (NEB, cat. #C2528) according to the manufacturer’s recommendations. Transformants were plated on LB + 100 μg/mL ampicillin (Amp) agar plates and incubated overnight growth at 37°C. Successful constructs were confirmed via Sanger sequencing and subsequent analysis in Geneious Prime (Auckland, New Zealand). Plasmid DNA from confirmed clones was purified using the GeneJet Plasmid Miniprep Kit (ThermoFisher Scientific; Waltham, MA; cat. #K0503) and transferred into *E. coli* SHuffle T7 express (NEB, cat. #C3029), T7 express lysY/lq (NEB, cat. #C3013), BL21(DE3) (NEB, cat. #C2527), BL21 (NEB, cat. #C2530), and NEB5α (NEB, cat. #C2987) via heat shock as described above.

**Table 4.**
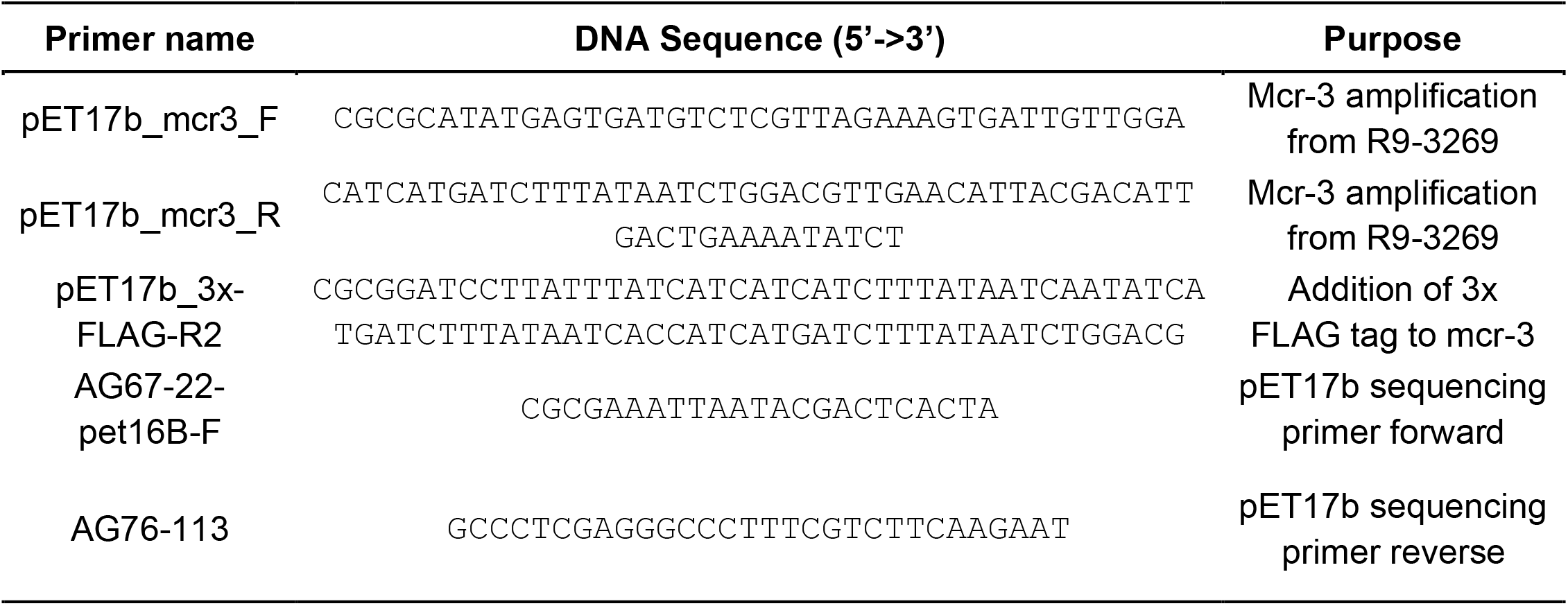
Primers used in this study

### Colistin susceptibility testing

Colistin susceptibility was tested by broth microdilution (BMD) according to the Clinical and Laboratory Standards Institute (CLSI) recommendations (61), except that PlateOne polypropylene 96-well plates (USAScientific; Ocala, FL; cat. #1837-9610) were used instead of polystyrene plates due to the known higher binding affinity of colistin to polystyrene (65, 66). Briefly, strains were grown in LB (+ 100 μg/mL Amp for pET17b plasmid carrying strains) at 37°C and 200 rpm shaking for 12-18 hours and subsequently diluted 1:200 fold in BBL Mueller-Hinton II broth cation-adjusted (MHII, BD cat. #212322) (+ 100 μg/mL Amp for pET17b plasmid carrying strains) followed by incubation at 37°C and 200 rpm shaking until cultures reached an OD600 of 0.3-0.4. Strains carrying pET17b plasmids were induced by the addition of 0.4 mM IPTG, 1 mM IPTG or filter-sterilized dH2O as a negative control and grown for another 2 hours. *E. coli* BL21(DE3) and T7 express lysY/l^q^ were induced at 37°C and 200 rpm shaking, while *E. coli* SHuffle T7 express was induced at 30°C, 200 rpm, based on the manufacturer’s recommendations (NEB). Strains were inoculated at a concentration of 5 x 10^5^ CFU/mL in MHII broth containing colistin (0.5 – 128 μg/mL) and IPTG (0, 0.4, or 1 mM) and grown statically for 16-20 hours at 35°C. *S. enterica* isolates R9-5409 and R9-3269 were used as controls, because R9-5409 does not encode any *mcr* variant and R9-3269 natively encodes *mcr-3;* their MIC values were previously reported as 0.5 μg/mL and 4 μg/mL, respectively.

### Alignment of *pmrA* and *pmrB* genes

The DNA sequences of *E. coli* str. K-12 substr. MG1655 genes *pmrA/basR* (locus tag: b4113) and *pmrB/basS* (locus tag: b4112) were downloaded from NCBI (Accession No. NC_000913.3). The *pmrA* and *pmrB* genes encoded by strains used in this paper were identified using BLAST (67), by querying the sequences against the genomes of NEB strains *E. coli* BL21 (Accession No. CP053601), BL21(DE3) (Accession No. CP053602), T7 express lysY/l^q^ (Accession No. CP053595), SHuffle T7 Express (Accession No. CP014269), and NEB5α (Accession No. CP017100.1). The identified *pmrA* and *pmrB* sequences were downloaded and converted into amino acid sequences using ExPASy, and both DNA and amino acid sequences were aligned using Clustal Omega (48, 68).

### Western Blot Analysis of MCR-3-FLAG

To confirm MCR-3*-FLAG* expression, *E. coli* B-strains containing the pET17b and pET17b-*mcr-3*-FLAG plasmid were grown at 37°C and 200 rpm shaking in LB + 100 g/mL Amp for 12-18 hrs, followed by 1:200 fold backdilution in MHII broth + 100 g/mL Amp until an OD600 of 0.3-0.4 was reached. Cells were induced with 0.4 mM IPTG or dH2O for 2 hrs and collected by centrifugation at 7,197 xg for 10 minutes (Sorvall X4RF PRO-MD centrifuge). Cells were resuspended in SDS-PAGE lysis buffer and lysed by 2 rounds of sonication using Branson Sonifier 50 sonicator (80% duty cycle, 7 output control) for 30 seconds on ice. Lysed cells were resolved on a 4% to 20% Mini-Protean TGX precast protein SDS-PAGE gel (Bio-Rad Laboratories; Hercules, CA, cat. #4568096). Resolved protein bands were visualized by activating the prestained SDS-PAGE gels with 1.5 min UV light exposure using the Bio-Rad ChemiDoc MP Imaging system. The proteins were transferred to polyvinylidene difluoride (PVDF) membrane from the Trans-Blot Turbo RTA Transfer Kit, PVDF (Bio-Rad Laboratories, cat. #170-4272) using the Trans-Blot Turbo transfer system (Bio-Rad Laboratories), according to the manufacturer’s recommendations. The membrane was blocked with TTBS (TBS buffer [50 mM Tris-Cl, 150 mM NaCl, pH 7.5] with 0.1% (v/v) Tween 20) containing 5% Blotting-Grade Blocker (Bio-Rad Laboratories, cat. #170-6404) for 30 mins, followed by incubation with the rabbit anti-FLAG primary antibody (Sigma, cat. #F7425, at 1:500 dilution) diluted in TTBS with 0.5% (W/V) skimmed milk powder at room temperature overnight with gentle shaking. The membrane was washed three times for 10 mins with TTBS before incubating with goat anti-rabbit-horseradish peroxidase secondary antibody (Thermo Fisher Scientific, cat. #65-6120, at 1:3,000 dilution) for 2 hours at room temperature with gentle shaking. The membrane was washed three times for 10 mins with TTBS and once for 10 mins with TBS before developing it with the Clarity Western ECL Substrate (Bio-Rad Laboratories, cat. #170-5061) and visualizing it using the Bio-Rad ChemiDoc MP Imaging system. Densitometric analysis of band intensity was carried out using the Bio-Rad Image Lab 6.1 software. Levels of MCR-3-FLAG expression were semi-quantitively estimated by normalizing the area under the curve (AUC) of the MCR-3-FLAG band on the developed PVDF membrane to the sum of th AUCs for all protein bands in the corresponding lane on the UV-activated SDS-PAGE gel (“total amount of protein”).

### Plating Assay

Cell viability in response to IPTG addition was tested by monitoring cell growth on agar plates. Briefly, strains were grown in LB broth (strains containing pET17b plasmids and constructs were supplemented with 100 μg/ml Amp) for 12-18 hours at 37°C and 200 rpm shaking. Strains were subsequently used as precultures to inoculate MHII broth tubes at 1:200-fold dilution (strains containing pET17b were supplemented with 100 μg/mL). MHII broth culture tubes were incubated at 37°C and 200 rpm shaking until an OD600 of 0.3 – 0.4 was reached. Ten-fold serial dilutions were prepared from each culture in phosphate-buffered saline, and 10 μl volumes were spotted onto Difco LB Miller (Luria-Bertani) Broth (BD, cat. #244620) agar plates (broth was supplemented with 15 g/L Bacto Agar [BD, cat. #214010]) supplemented with 0 mM, 0.4 mM or 1 mM of IPTG and/or 1 mM MgCl_2_. Plates were incubated at 37°C overnight. Pictures were taken with the Bio-Rad ChemiDoc MP imaging system. Cell viability was determined by monitoring the highest dilutions with visible colonies and inspecting colony size and morphology.

## Acknowledgements

We want to thank Bailey Lubinski for the construction of the pET17b-mcr-3-FLAG expression plasmid.

This work is supported by a Hatch grant under accession number 1023966 from the USDA National Institute of Food and Agriculture. The funders had no role in study design, data collection and interpretation, or the decision to submit the work for publication

